# SARS-CoV-2 Triggers Complement Activation through Interactions with Heparan Sulfate

**DOI:** 10.1101/2022.01.11.475820

**Authors:** Martin W. Lo, Alberto A. Amarilla, John D. Lee, Eduardo A. Albornoz, Naphak Modhiran, Richard J. Clark, Vito Ferro, Mohit Chhabra, Alexander A. Khromykh, Daniel Watterson, Trent M. Woodruff

## Abstract

The complement system has been heavily implicated in severe COVID-19 with clinical studies revealing widespread gene induction, deposition, and activation. However, the mechanism by which complement is activated in this disease remains incompletely understood. Herein we examined the relationship between SARS-CoV-2 and complement by inoculating the virus in lepirudin-anticoagulated human blood. This caused progressive C5a production after 30 minutes and 24 hours, which was blocked entirely by inhibitors for factor B, C3, C5, and heparan sulfate. However, this phenomenon could not be replicated in cell-free plasma, highlighting the requirement for cell surface deposition of complement and interactions with heparan sulfate. Additional functional analysis revealed that complement-dependent granulocyte and monocyte activation was delayed. Indeed, C5aR1 internalisation and CD11b upregulation on these cells only occurred after 24 hours. Thus, SARS-CoV-2 is a non-canonical complement activator that triggers the alternative pathway through interactions with heparan sulfate.

## INTRODUCTION

COVID-19 is a highly contagious respiratory infection caused by the severe acute respiratory syndrome coronavirus 2 (SARS-CoV-2). In the last two years, this disease has affected over 300 million individuals and caused over 5.4 million deaths (1). Thus, unprecedented efforts have been put towards vaccine and drug development, but with the possibility of new variants and the inevitability of future pandemics, a fundamental understanding of severe COVID-19 is still needed. In this context, SARS-CoV-2 replicates in an unchecked manner and evades the immune system by exploiting several inborn and acquired weaknesses (2, 3). At a critical mass, these virions then trigger a hyperinflammatory response that results in acute respiratory distress syndrome (ARDS) (4). Emerging evidence suggests that the complement system plays a key role in this process (5-7). Indeed, complement activation has been correlated with disease severity (8) and small case studies have shown that complement inhibition can be effective in critical patients, prompting at least six anti-complement drugs to be taken to clinical trials (5). However, whilst *in vitro* mechanistic studies have demonstrated that specific viral proteins can activate complement, the relationship between SARS-CoV-2 and complement activation remains incompletely understood.

Complement-mediated disease in COVID-19 appears to be confined to severely ill patients who are unable to bring the virus under immunological control. In these patients, SARS-CoV-2 exploits defects in the type 1 interferon system and replicates in an unchecked manner, which at a critical mass, is believed to drive a form of complement-mediated hyperinflammation (3). Indeed, evidence of complement activation has been correlated with disease severity and includes serum C5a and C5b-9 concentrations (8), monocyte and granulocyte CD11b expression (9) which can be due to C5aR1 activation (10, 11), and post-mortem immunochemistry (7, 12). These features occur on the background of airway and intravascular complement synthesis (13, 14) and are particularly prominent in individuals who are genetically prone to C5 cleavage (15), who have elevated mannose binding protein levels (16), or who have reduced CD55 expression (17). Additional investigations suggest that complement activation in severe COVID-19 can occur through the classical, lectin, and alternative pathways (18-20). Thus, complement is likely to be a key driver of severe COVID-19.

Moreover, molecular investigations have provided some insight into the underlying mechanisms that drive complement activation in COVID-19. Indeed, an initial study utilising the SARS-CoV-2 S-protein in a specialised functional assay suggests that the virus may activate the alternative pathway by binding to cell surface heparan sulfate, which disinhibits factor H-mediated complement suppression (21). In this study, normal human serum pre-treated with recombinant SARS-CoV-2 S protein caused complement deposition and cytotoxicity in complement-inhibitor deficient cells. However, SARS-CoV-2 S-protein did not generate complement activation products in human serum without such cells or after heparan sulfate or factor H supplementation. In addition, a more recent study found that the SARS-CoV-2 S and N proteins are able to activate the lectin pathway via MASP-2 (22). Thus, early molecular studies using viral proteins suggest that SARS-CoV-2 can directly activate the complement system, but conclusive evidence for this with live virus is still outstanding.

Therefore, here we inoculated SARS-CoV-2 into lepirudin-anticoagulated human blood and used ELISAs and flow cytometry to measure complement activation and functionality respectively. We show that SARS-CoV-2 activates complement via the alternative pathway by interacting with heparan sulfate, and in doing so causes delayed leukocyte activation through C5a-C5aR1 signalling.

## METHODS

### Study approval

This research was approved by the University of Queensland Human Research Ethics Committee and the University of Queensland Biosafety Committee (supplementary methods 1). All participants gave informed written consent.

### Participants

Whole blood was drawn from healthy individuals (supplementary table 1) with a 4.9ml S-Monovette (SARSTEDT, Nümbrecht, Germany, # 04.1926.001) and anticoagulated with 50μg/ml Lepirudin (Pharmion, Boulder, California), which is an anticoagulant that permits *ex vivo* complement activation (11). Türk’s solution (Sigma-Aldrich, Saint Louis, Missouri, # 109277) was used as per manufacturer guidelines to perform total leukocyte counts for MOI calculations.

### Whole blood inoculation

For the ELISA experiments, 100μl of whole blood was inoculated with SARS-CoV-2 at a MOI of 0.1 or 1.0 (supplementary methods 2), LPS O111:B4 (200μg/ml; Sigma-Aldrich, #L2630-100MG), or a mock solution (i.e., DMEM with 2% HIFCS and P/S) for 30 minutes or 24 hours at 37°C. Samples were then centrifuged at 2000*g* for 10 minutes at 4ºC and plasma was aliquoted and stored at -80ºC for downstream analysis. Certain samples were pre-treated with the following for 30 minutes at 37°C: SFMI-1 (MASP1/2 inhibitor 10μM; synthesized in house (23)), LNP023 (factor B inhibitor, 10μM; AdooQ Bioscience, Irvine, California, #A18905) compstatin analogue (C3 inhibitor, 20μM; Wuxi AppTec Ltd, Shanghai, China, #C15031904), eculizumab (C5 inhibitor, 100μg/mL; Ichorbio, Wantage, United Kingdom, #ICH4005)), EGCG (heparan sulfate inhibitor, 100μM; Sigma-Aldrich E4143-50MG), or pixatimod/PG545 (heparan sulfate mimetic, 100μg/mL; synthesized in house (24)). This assay was repeated with plasma isolated from whole blood after centrifugation at 2000*g* for 10 minutes at room temperature. For the flow cytometry experiments, whole blood was mixed 1:1 with RPMI1640 (Gibco, Waltham, Massachusetts, #42401-018) and inoculated with SARS-CoV-2 at a MOI of 0.1 or 1.0 or a mock solution (as above) and incubated for 3 or 24 hours at 37°C with 5% CO_2_. Certain samples were pre-treated with PMX205 (10μM; synthesized as previously described (25)) or eculizumab (as above) for 30 minutes at 37°C with 5% CO_2_.

### ELISA

A C5a ELISA (R&D, Minneapolis, Minnesota, #DY2037) was performed as per manufacturer’s guidelines on plasma samples from whole blood inoculated with SARS-CoV-2 for 30 minutes and 24 hours.

### Flow cytometry

Whole blood inoculated with SARS-CoV-2 was blocked with 5μl of TruStain (Biolegend, San Diego, California, # 422302) for 10 minutes at room temperature and then stained for granulocyte and monocyte markers, complement receptors, and viability for 15 minutes at room temperature (supplementary methods 3). Samples were then fixed and lysed with 2ml of BD FACSLyse (BD, Franklin Lakes, New Jersey, # 349202) for 15 minutes at room temperature and inverted 10 times at the 0-and 7.5-minute marks. Lysed samples were then centrifuged at 600*g* for 5 minutes at room temperature. Leukocytes were then resuspended in PBS for flow cytometry acquisition on a BD LSR Fortessa II. Data analysis was performed with FlowJo v10.6.2.

### Statistical analysis

Statistical analysis was performed with GraphPad Prism Software v9.3.1. A one-way ANOVA with Dunnett’s post-test analysis was used for one factor ordinal and categorical data. Otherwise, t-tests were used to compare means to assess for temporal differences in functional assays and for drug effects in the context of SARS-CoV-2 inoculation. Additional detail is provided in supplementary methods 4 and the source data file.

## RESULTS AND DISCUSSION

### SARS-CoV-2 activates the alternative pathway through interactions with heparan sulfate

We first investigated whether SARS-CoV-2 could mediate complement activation by inoculating the virus at multiplicity of infection (MOI) 0.1 and 1.0 in lepirudin-anticoagulated human blood from individuals with no history of COVID-19 or respective vaccination and subsequently measuring plasma C5a with an ELISA. Indeed, inoculated whole blood showed an increase in C5a of 5-10ng/mL and 30-50ng/mL at MOI 1.0 when compared to the mock control at 30 minutes and 24 hours post-inoculation respectively (figure 1a). However, this phenomenon could not be replicated in isolated plasma (figure 1b), which suggests that a cellular component is required. To further delineate the pathways involved, SARS-CoV-2 inoculated whole blood was pre-treated with inhibitors/mimetics, in which antagonism of factor B, C3, C5, and heparan sulfate, and to a lesser extent MASP1/2, attenuated complement activation (figure 2).

**Figure 1.**
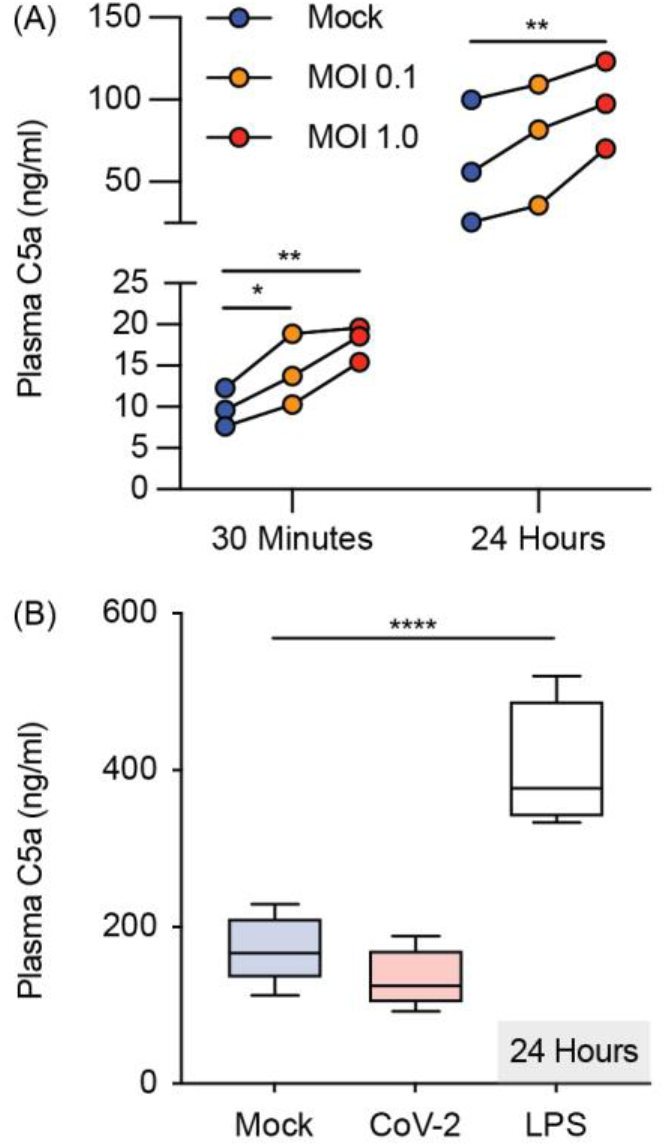
SARS-CoV-2 activates complement in whole blood but not in plasma. Plasma C5a levels were assessed with an ELISA in SARS-CoV-2 inoculated lepirudin-anticoagulated (A) human blood (n = 3) and (B) plasma at indicated times (n = 5). Boxes depict medians and inter-quartile ranges and have Tukey whiskers. MOI = Multiplicity of infection; CoV-2 = SARS-CoV-2 MOI 1.0; * P<0.05, ** P<0.01, **** P<0.0001, using a one-way ANOVA and Dunnett’s post-test.

**Figure 2.**
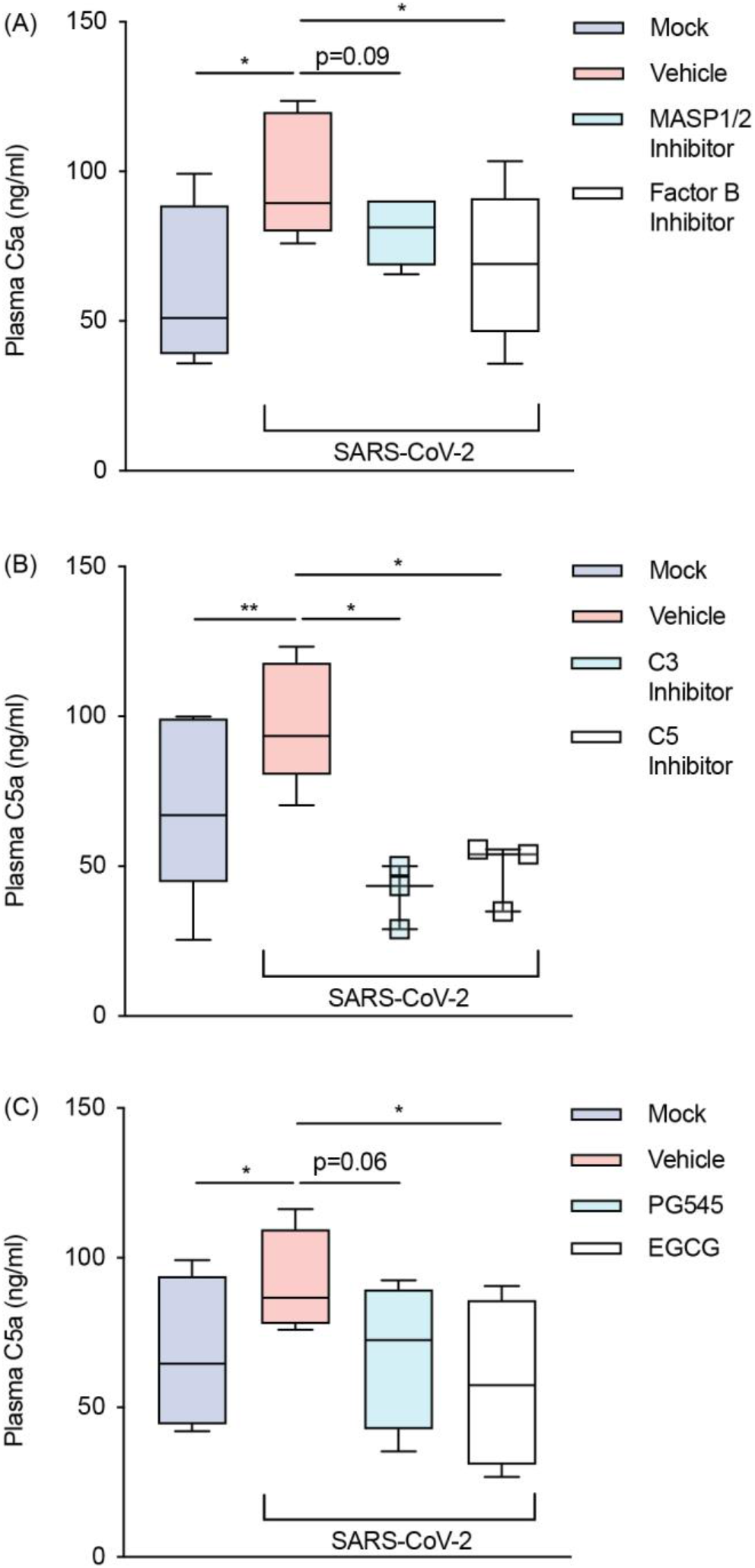
SARS-CoV-2 activates the alternative pathway through interactions with heparan sulfate. Plasma C5a levels were assessed with an ELISA in SARS-CoV-2 inoculated lepirudin-anticoagulated human blood pre-treated with various inhibitors and mimetics. These included antagonists for (A) MASP1/2 (SFMI-1 10μM) and factor B (LNP023 10μM) (n = 5); (B) C3 (Compstatin 20μM) and C5 (Eculizumab 100μg/mL) (n = 3); and (C) heparan sulfate (EGCG 100μM) as well as a mimetic for heparan sulfate (PG545 100μg/mL) (n = 4). SARS-CoV-2 was inoculated at MOI 1.0 for 24 hours. Boxes depict medians and inter-quartile ranges and have Tukey whiskers. Where this was not possible, individual values are presented instead. MOI = multiplicity of infection; * P<0.05, ** P<0.01, using a paired one-tailed t test. All blood donors were seronegative for anti-SARS-CoV-2 antibodies (Figure 2 - figure supplement 1).

Moreover, serological testing confirmed that all blood donors were seronegative for SARS-CoV-2 and thus excluded any antibody-mediated complement activation via the classical pathway (figure 2 – figure supplement 1). Thus, given that the C3, C5, and heparan sulfate inhibitors were membrane impermeable, these findings indicate that SARS-CoV-2 activates the alternative pathway through interactions with cell surface heparan sulfate and subsequent plasma complement deposition.

### Flow cytometry optimization and incidental findings on monocyte activation

Next, we sought to determine if SARS-CoV-2-mediated complement activation was sufficient to induce a functional response. As C5a stimulation of myeloid cells causes C5aR1 internalisation and CD11b upregulation at the cell surface (10, 11), we inoculated whole blood with SARS-CoV-2 (MOI 0.1 and 1.0) for 3 and 24 hours and used flow cytometry to measure C5aR1 and CD11b surface expression on neutrophils, eosinophils, and monocytes. To do so, we trialled an optimized flow cytometry panel under inoculation conditions and found gating properties to be slightly altered (figure 3 – figure supplement 1) compared to blood inoculated with a mock control. Indeed, whilst granulocyte markers were unaffected, monocyte markers including CD16 and HLA-DR were upregulated in an MOI-dependent fashion (supplementary figure 2a), which suggests that other immune phenomena were present in this assay. Thus, pan-monocytes without subset differentiation were examined in this study.

### SARS-CoV-2 causes delayed complement-mediated leukocyte activation

Given that immune stimulators induce rapid changes in myeloid cell activation, we first tested whether SARS-CoV-2 could cause complement-mediated leukocyte activation at 3 hours post-inoculation in whole blood. However, at this time point, flow cytometry revealed few alterations in cell activation markers in neutrophils, eosinophils, and monocytes. We therefore extended the period of SARS-CoV-2 incubation to 24 hours and detected significant C5aR1 internalisation and CD11b upregulation on innate leukocytes. This was observed in neutrophils, eosinophils, and monocytes and was most prominent at MOI 1.0 (figure 3). Interestingly, at 3 hours post-inoculation, CD11b upregulation without concomitant C5aR1 internalisation was noted in neutrophils, which suggests that these cells can react acutely to SARS-CoV-2 through receptors independent of complement (figure 3 – figure supplement 2).

**Figure 3.**
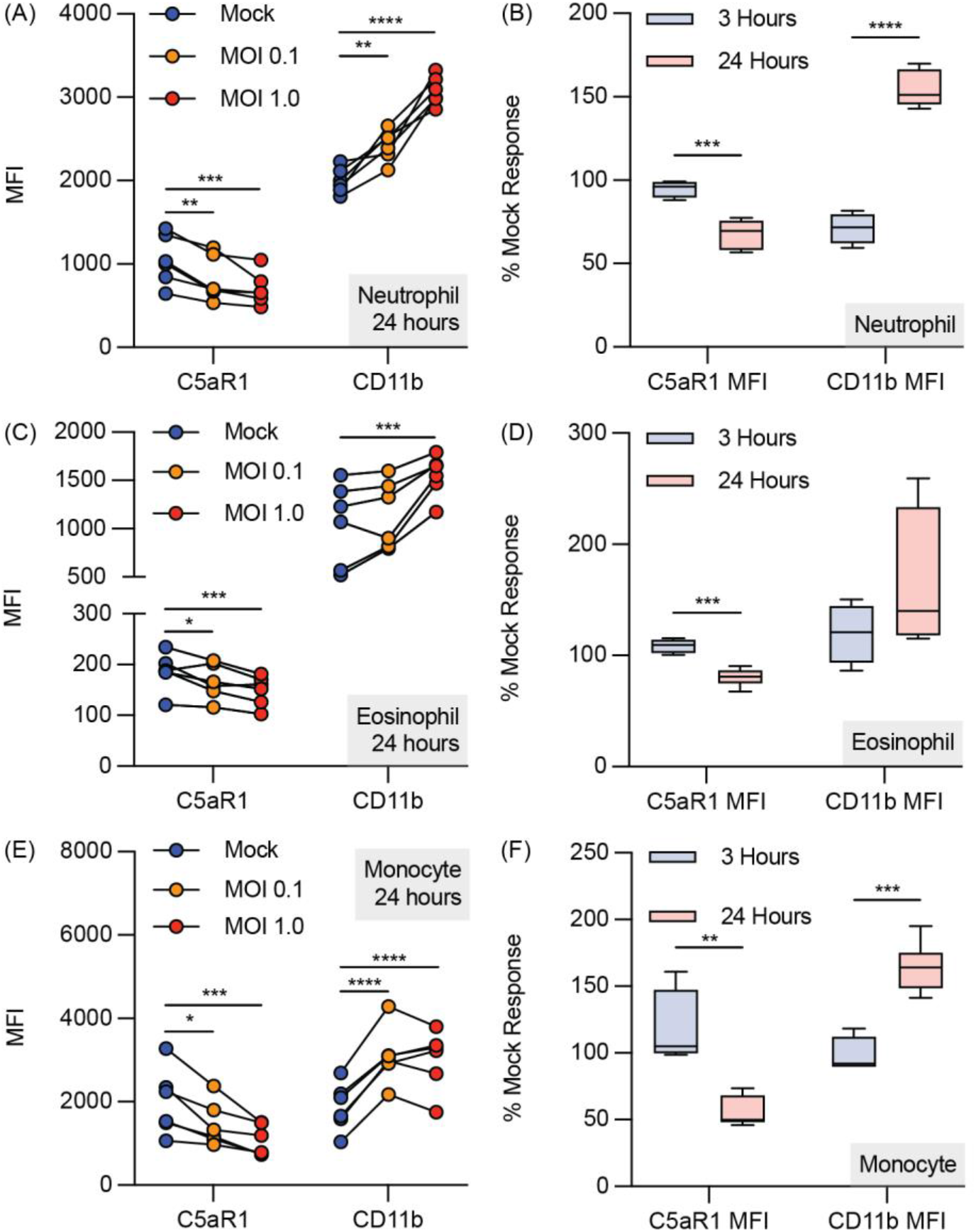
SARS-CoV-2 causes delayed complement-mediated activation in innate leukocytes. SARS-CoV-2 inoculated lepirudin-anticoagulated human blood was analysed with flow cytometry. SARS-CoV-2 caused dose-dependent C5aR1 internalisation and CD11b upregulation in (A) neutrophils, (C) eosinophils, and (E) monocytes at 24 hours post inoculation (n = 6). Comparison of C5aR1 and CD11b responses between SARS-CoV-2 inoculated blood at 3 (n = 4) and 24h (n = 6) expressed as a percentage change from mock inoculation (B, D, F). Boxes depict medians and inter-quartile ranges and have Tukey whiskers. MOI = multiplicity of infection; * P<0.05, ** P<0.01, *** P<0.001, **** P<0.0001 using a one-way ANOVA and Dunnett’s post-test or unpaired t test with two stage step-up 1% false discovery rate correction. Flow cytometry gating strategy and incidental findings are provided (Figure 3 – figure supplement 1 and 2).

### Anti-C5/C5aR1 drugs eculizumab and PMX205 inhibit SARS-CoV-2-induced complement-mediated inflammation

To investigate the functional role of the terminal complement pathway in mediating leukocyte activation in response to SARS-CoV-2, we next pre-incubated whole blood with a C5 inhibitor (eculizumab) and a C5aR1 antagonist (PMX205). At 24 hours post inoculation, both drugs inhibited SARS-CoV-2-dependent C5aR1 internalisation and CD11b upregulation in neutrophils and eosinophils with similar efficacy (figure 4a-b). By contrast, on monocytes, these drugs were only able to partially inhibit C5aR1 internalisation and did not lower CD11b upregulation (figure 4b). This latter finding suggests that complement at the level of C5 is not involved in this process in monocytes.

**Figure 4.**
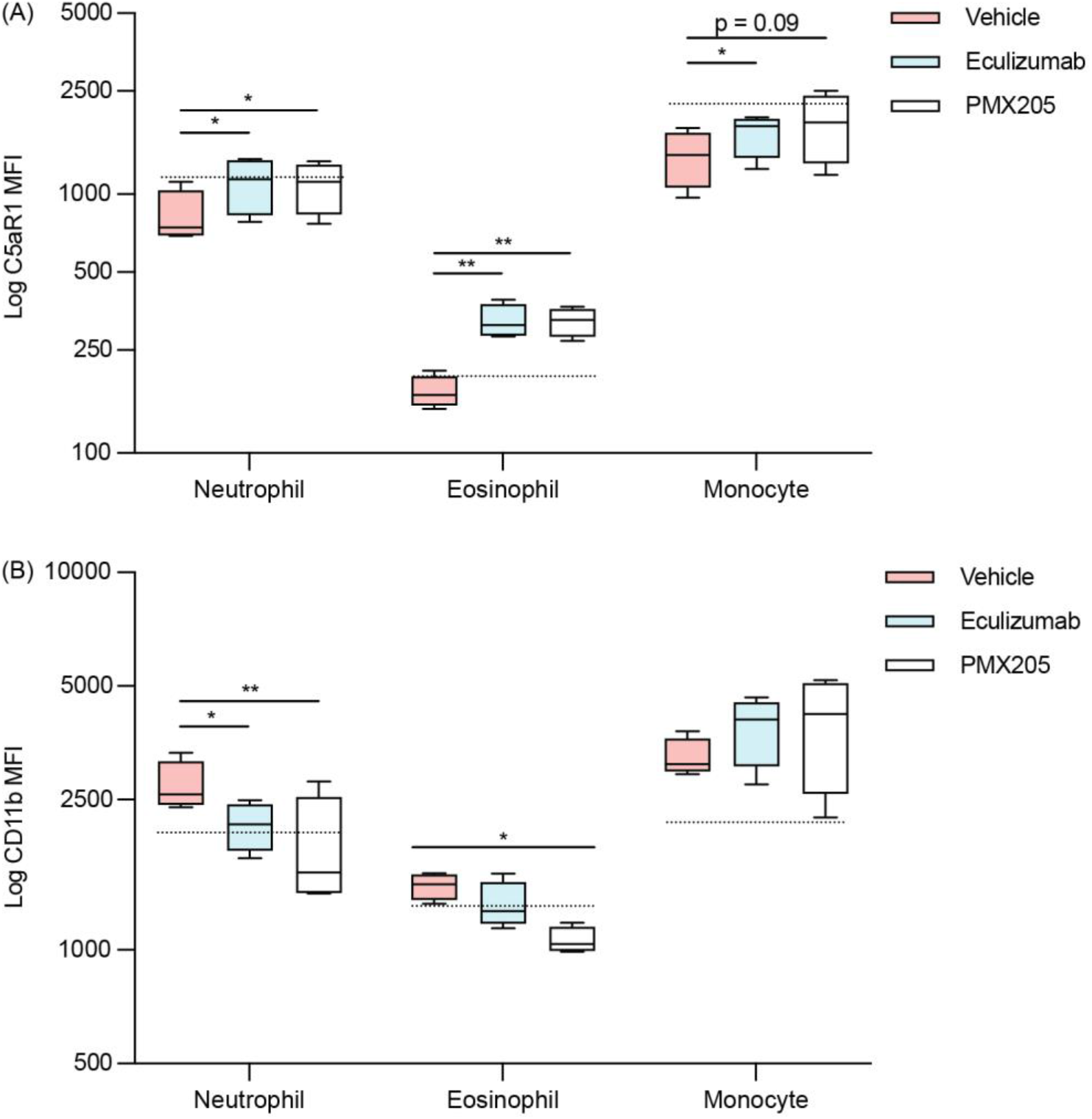
C5/C5aR1 inhibition attenuates SARS-CoV-2-induced leukocyte activation. Anti-C5/C5aR1 inhibitors eculizumab (100μg/mL) and PMX205 (10μM) were administered to lepirudin-anticoagulated whole blood (n = 4) prior to SARS-CoV-2 inoculation at MOI 1.0 for 24 hours and analysed with flow cytometry. Activation markers C5aR1 and CD11b were measured as MFI. Boxes depict medians and inter-quartile ranges and have Tukey whiskers. The dashed line represents the mock infection baseline. MOI = multiplicity of infection; MFI = median fluorescence intensity; * P<0.05, ** P<0.01, using a paired one-tailed t test.

### Results support clinical findings of complement in COVID-19

Overall, the complement activation observed in our *ex-vivo* blood study is consistent with the current clinical and molecular understanding of COVID-19. Foremostly, *in vivo* complement profiling in severe COVID-19 reveals hyperactivation of the alternative pathway (14, 19) and *in vitro* studies have shown that viral S-protein can activate this same pathway through interactions with heparan sulfate (21). However, in contrast to our *ex vivo* model, complement activation during COVID-19 is probably a multifactorial phenomenon driven by multiple complement pathways (7, 20, 22), non-specific DAMP release (5), genetic susceptibility to complement activation (15-17), and local complement synthesis (13). But in this regard, our participants were seronegative for anti-SARS-CoV-2 antibodies, which would imply that the classical pathway was not activated in this study. Additionally, given that C5a-mediated immune activation typically occurs within 60 minutes (11, 26), the delayed response requiring 24 hours in our model is consistent with the gradual progression (4) and upregulation of CD11b on leukocytes in severe cases (9). Thus, these results strongly support the emerging paradigm of heparan sulfate- and alternative pathway-mediated disease in severe COVID-19.

### Complement activation appears to be localised to organs that support replication

When placed in the context of *in vivo* viral titres, our results also suggest that SARS-CoV-2-mediated complement activation is localized to organs that can support replication. On average COVID-19 patients have median viral titres of ∼10^6^ RNA copies/ml in their airways; as determined from nasopharyngeal swaps, sputum, saliva, and bronchoalveolar lavage fluid; and ∼10^3^ RNA copies/ml in their serum, in which the former is at least 10 times higher in severe cases compared to that in mild cases (27, 28). By comparison, in this study whole blood inoculated with SARS-CoV-2 at MOI 0.1 and 1.0 was exposed to virion concentrations of ∼3.5-5.5 × 10^5^ and ∼3.5-5.5 × 10^6^ FFU/ml respectively. Thus, complement activation in severe COVID-19 is most likely localised to tissues that support replication (e.g., lung parenchyma) with changes in plasma complement occurring as a secondary phenomenon. This implies that anti-complement drugs in severe COVID-19 require significant tissue distribution/permeation, which may have been a limiting factor for the parenteral drugs that have so far been tested in clinical trials. In this sense, SARS-CoV-2 can be viewed as an organ-based activator of complement that poses unique challenges to drug development.

### Complement and heparan sulfate in severe COVID-19

Furthermore, complement activation through heparan sulfate appears to be an inadvertent phenomenon. Indeed, this polysaccharide post-translational modification promotes specific protein-protein interactions, which in this case concentrates factor H on host cells to prevent excessive complement activation (29). Therefore, whilst SARS-CoV-2 principally uses heparan sulfate to dock with angiotensin-converting enzyme 2 to enable infection (30), it also secondarily activates complement by causing factor H disinhibition. In our *ex vivo* model, this was largely mediated by CD44v3 on lymphocytes, but *in vivo* is probably the result of interactions with a range of membrane bound (e.g., syndecan 1-4 and glypican 1-6) and extracellular matrix proteoglycans (e.g., perlecan, agrin, and collagen XVII) (31). Interestingly, S protein can bind to heparan sulfate and disrupt anti-thrombin and heparin cofactor II activity and thus this polysaccharide modification warrants further investigation as a multi-faceted drug target in severe COVID-19 (32).

### Concluding remarks

COVID-19 continues to cause more deaths than any other pandemic in living memory. Mounting evidence suggests that complement plays a key role in its most severe form and here we show that SARS-CoV-2 interacts with heparan sulfate to activate the alternative pathway, which ultimately drives innate leukocyte activation through C5a-C5aR1 signalling. In doing so, these findings support the use of targeted anti-complement treatments in severe COVID-19.

## AUTHORSHIP CONTRIBUTIONS

MWL* designed the project, recruited healthy participants, managed logistics, ran the flow cytometry samples, analysed the data, made the figures, and drafted the manuscript.

AAA* contributed to experimental design, conducted all experimental work within the biosafety level 3 facility, and drafted the methods section for this.

JDL contributed ideas, validated reagents, and edited the final manuscript. EAA edited the final manuscript and assisted AAA in experimental planning. NM performed serology experiments and drafted the methods section for this. RJC synthesized SFMI-1.

MC synthesized Pixatimod/PG545 under the supervision of VF.

AAK oversaw SARS-CoV-2 laboratory setup and assay development, provided funding and edited the final manuscript.

DW oversaw the SARS-CoV-2 experimental studies, provided funding, and edited the final manuscript.

TMW co-conceived the idea for the project, contributed to experimental design, oversaw the entire study, provided funding, helped design figures, and contributed substantial edits to the drafted manuscript.

The order of co-first authorship* was based on the intellectual contribution to the manuscript.

## ACKNOWLEDGMENTS

We would like to thank the Queensland Health Forensic and Scientific Services, Queensland Department of Health, for providing the QLD02 SARS-CoV-2 isolate. We acknowledge funding support from the National Health and Medical Research Council (APP1118881), The Australian Infectious Diseases Research Centre (COVID-19 seed grant to AAK), and the Medical Research Future Fund (APP1202445-2020 MRFF Novel Coronavirus Vaccine Development Grant).

## COMPETING INTERESTS

Trent M. Woodruff is an inventor on patents pertaining to complement inhibitors for inflammatory diseases. He has consulted to Alsonex Pty Ltd (who are commercially developing PMX205) and has received honorarium from Alexion Pharmaceuticals (who developed eculizumab) for participation in industry conferences and meetings. Vito Ferro is an inventor on patents for Pixatimod/PG545. All other authors declare no conflicts of interest.

## SUPPLEMENTARY MATERIAL

**Supplementary Table 1:** Research Participants for this Study. Participants were recruited from the local Brisbane area and had no history of COVID-19, no history of acute illness or vaccination in the last 2 weeks, no immunodeficiencies or autoinflammatory/autoimmune conditions, and were not on any immunomodulatory medications (e.g., corticosteroids). No significant sex or age differences were found between the cohorts at different time points for the ELISA and flow cytometry experiments.

| C5a ELISA 30min |  | C5a ELISA 24h |  | C5a ELISA Pathways |  |
| --- | --- | --- | --- | --- | --- |
| *Sex* | *Age (Years)* | *Sex* | *Age (Years)* | *Sex* | *Age (Years)* |
| Male | 30 | Male | 40 | Female | 20 |
| Male | 23 | Male | 26 | Female | 20 |
| Female | 21 | Male | 36 | Male | 31 |
|  |  |  |  | Male | 27 |
|  |  |  |  | Female | 57 |
| Flow Cytometry 3h |  | Flow Cytometry 24h |  |  |  |
| *Sex* | *Age (Years)* | *Sex* | *Age (Years)* |  |  |
| Male | 36 | Male | 32 |  |  |
| Male | 41 | Male | 24 |  |  |
| Male | 25 | Male | 30 |  |  |
| Female | 26 | Female | 19 |  |  |
|  |  | Male | 23 |  |  |
|  |  | Female | 25 |  |  |

**Figure 2 – figure supplement 1.**
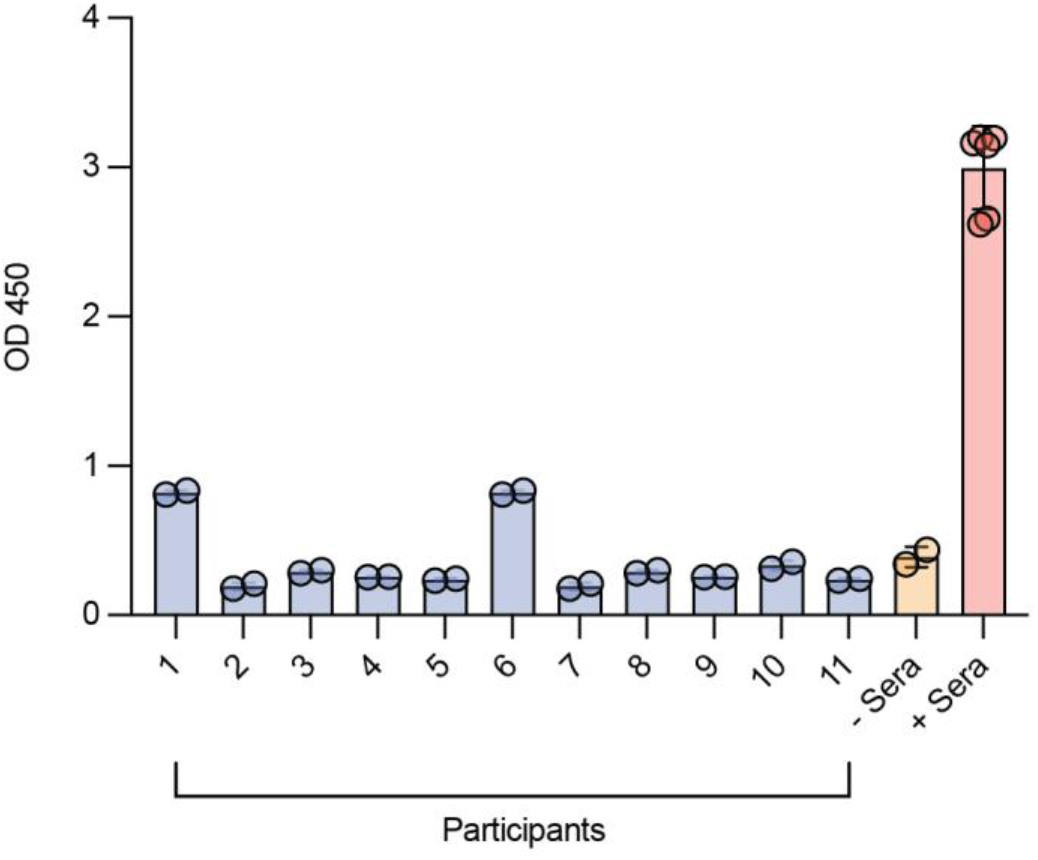
SARS-CoV-2 serology testing of plasma from whole blood used for C5a ELISA studies. Pre-COVID-19 serum (-Sera) was used as a negative control and a biological standard NIBSC 20/130 (+ Sera) was used as a positive control. The data here are from samples diluted at 1:10. Full methods are provided in supplementary methods 5; OD 450 = optical density at the wavelength of 450nm

**Figure 3 – figure supplement 1.**
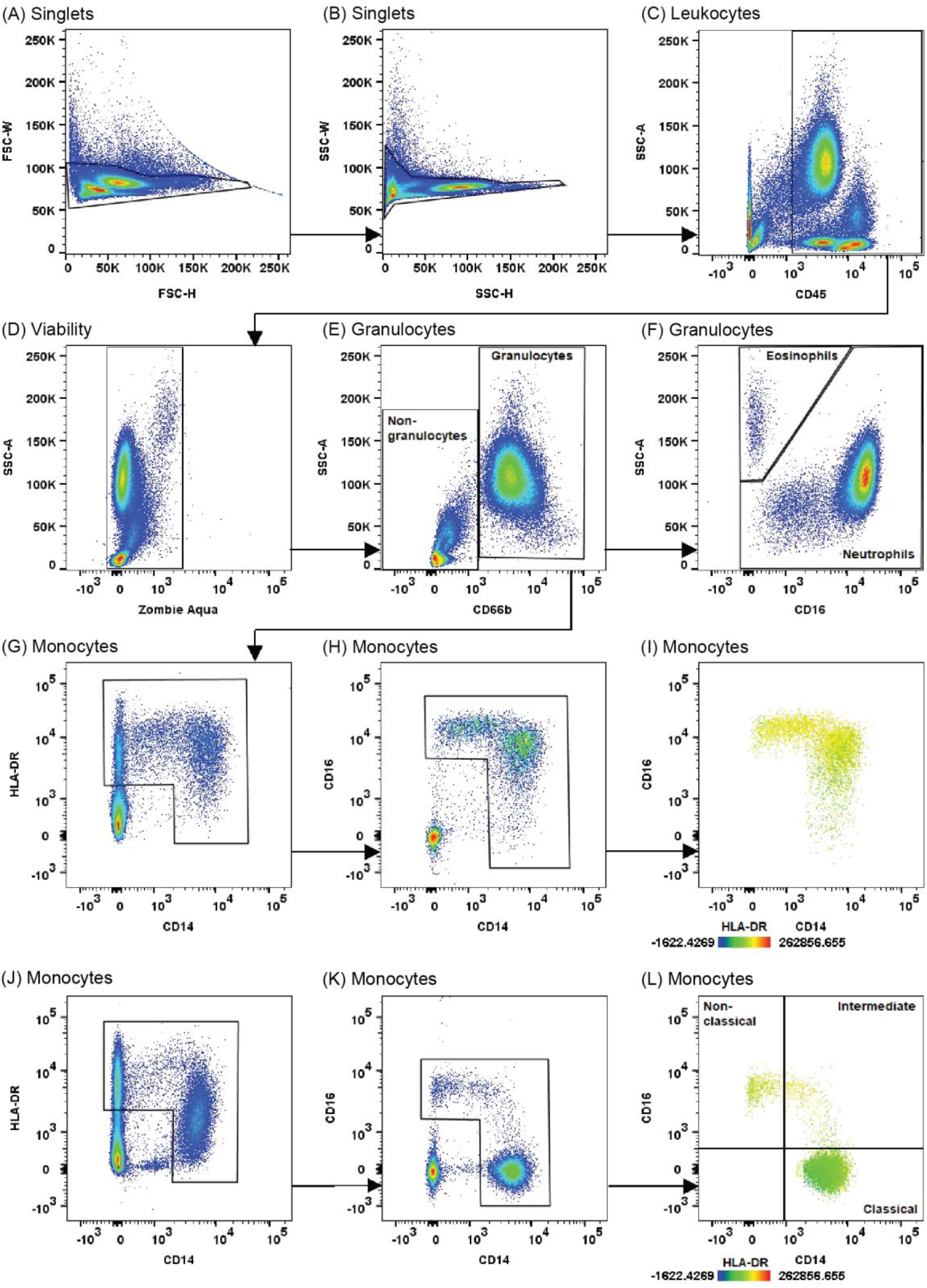
Flow cytometry gating strategy. Representative plots of SARS-CoV-2 inoculated whole blood stained with fluorophore-antibodies and a viability dye for flow cytometry (A-I). For comparison, representative plots of virus-naïve whole blood stained in the same fashion are also provided for the monocyte gates (J-L). All samples had >95% leukocyte viability.

**Figure 3 – figure supplement 2.**
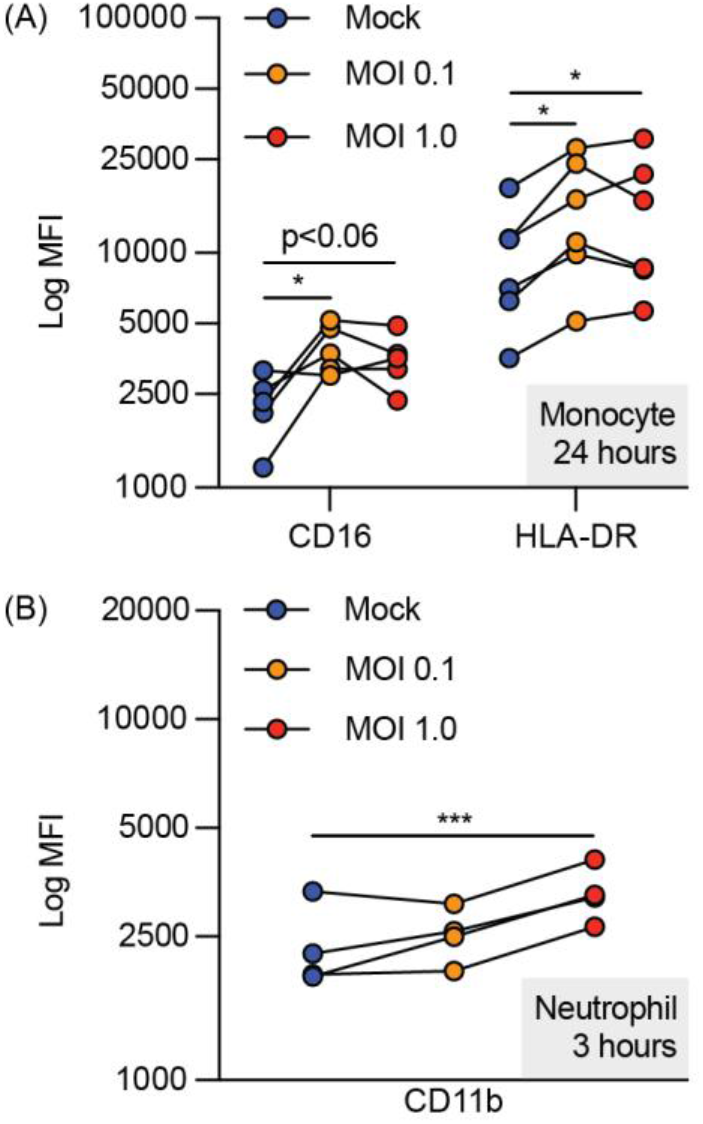
Upregulation of (A) CD16 and HLA-DR on monocytes and (B) CD11b on neutrophils exposed to SARS-CoV-2. SARS-CoV-2 inoculated lepirudin-anticoagulated whole blood was analysed with flow cytometry at 24 (n = 5-6) and 3 hours post-inoculation (n = 4) respectively. Surface markers were quantified as MFI. MOI = multiplicity of infection; MFI = median fluorescence intensity; * P<0.05, *** P<0.001 using a one-way ANOVA and Dunnett’s post-test.

**Supplementary Methods 1: Institutional Approval**

This research was approved by the University of Queensland Human Research Ethics Committee (TW00105) and the University of Queensland Biosafety Committee (IBC/390B/SCMB/2020, IBC/1301/SCMB/2020, IBC/376B/SBMS/2020 and IBC/447B/SCMB/2021). All experiments were performed in the biosafety level 3 facility at the School of Chemistry and Molecular Biosciences at The University of Queensland, Australia.

**Supplementary Methods 2: Cell Line and Virus**

An African green monkey kidney cell line (Vero E6 cells) was cultured in DMEM (Gibco) supplemented with 10% heat inactivated foetal calf serum (HIFCS) (Bovogen, Melbourne, Australia) and 100U/mL of penicillin and 100μg/mL of streptomycin (P/S). Cells were maintained at 37 ºC with 5 % CO_2_. The Queensland SARS-CoV-2 isolate QLD02 (GISAID accession EPI_ISL_407896) was recovered from a patient on 30/01/2020 by the Queensland Health Forensic & Scientific Services. VeroE6 cells were then inoculated with this isolate and an aliquot (passage 2) was provided. Viral stock (passage 3) was then generated through inoculation of VeroE6 cells in DMEM with 2% HIFCS and P/S and stored at -80 ºC. The viral titre was determined by an immuno-plaque assay (iPA) as previously described (33).

**Supplementary Methods 3: Flow Cytometry Staining Reagents**

Reagents included CD14-PerCP-Cy5.5 (Biolegend, # 301824), CD16 APC-Cy7 (Biolegend, # 302018), HLA-DR BV785 (Biolegend, # 307642), CD88 (C5aR1) PE-Cy7 (Biolegend, # 344308), CD45 BV605 (Biolegend, # 368524), CD66b FITC (Biolegend, # 305104), CD11b AF700 (Biolegend, # 301356), and Zombie Aqua (Biolegend, #423102).

**Supplementary Methods 4: Statistical Analysis**

Initially, a sample size of 3 was determined to be most appropriate for this study. This was decided in reference to previous *ex vivo* complement experiments that have been conducted with the lepirudin whole blood system (34). However, we also generated additional biological replicates according to donor availability and achieved final sample sizes of between 3 and 6.

Each experiment was performed on one occasion and all data reflects biological replication except for that in supplementary figure 3, in which data points reflect technical replication. Biological and technical replication was defined as the replication of an assay in a blood sample from a distinct or the same donor respectively. One outlier was excluded in supplementary figure 2a as its mock data point was more than 3 standard deviations from the mean. No other data was excluded.

**Supplementary Methods 5: SARS-CoV-2 Serology Testing**

Trimeric SARS-CoV-2 spike protein was coated at 2 μg/ml on an ELISA plate overnight (35). Plates were blocked for 1 hour at room temperature with a blocking buffer (PBS containing 0.05% Tween-20 and milk sera diluent/blocking solution (Seracare, Milford, Massachusetts)). Plasma from the whole blood used for the C5a ELISA study, a positive plasma control NIBSC 20/130 and a pre-COVID-19 serum negative control were serially diluted in blocking buffer and added to the plate for 1 hour at 37 °C. Plates were washed and probed by goat anti-human HRP antibody (1:2500) in blocking buffer for 1 hour in 37 °C. Tetramethylbenzidine substrate solution and sulfuric acid stop solution were then added prior to absorbance analysis. NIBSC 20/130A is a human covalence serum obtained from National Institute for Biological Standards and Control (URL: https://www.nibsc.org/documents/ifu/20-130.pdf).

## REFERENCES

1. Dong E, Du H, Gardner L. An interactive web-based dashboard to track COVID-19 in real time. The Lancet Infectious Diseases. 2020;20(5):533–4.

2. Taefehshokr N, Taefehshokr S, Hemmat N, Heit B. Covid-19: Perspectives on Innate Immune Evasion. Frontiers in immunology. 2020;11(2549).

3. Hadjadj J, Yatim N, Barnabei L, Corneau A, Boussier J, Smith N, et al. Impaired type I interferon activity and inflammatory responses in severe COVID-19 patients. Science (New York, NY). 2020;369(6504):718–24.

4. Berlin DA, Gulick RM, Martinez FJ. Severe Covid-19. New England Journal of Medicine. 2020.

5. Lo MW, Kemper C, Woodruff TM. COVID-19: Complement, Coagulation, and Collateral Damage. The Journal of Immunology. 2020:ji2000644.

6. Woodruff TM, Shukla AK. The Complement C5a-C5aR1 GPCR Axis in COVID-19 Therapeutics. Trends in immunology. 2020;41(11):965–7.

7. Satyam A, Tsokos MG, Brook OR, Hecht JL, Moulton VR, Tsokos GC. Activation of classical and alternative complement pathways in the pathogenesis of lung injury in COVID-19. Clinical Immunology. 2021:108716.

8. Ma L, Sahu SK, Cano M, Kuppuswamy V, Bajwa J, McPhatter JN, et al. Increased complement activation is a distinctive feature of severe SARS-CoV-2 infection. Science Immunology. 2021;6(59):eabh2259.

9. Gupta R, Gant VA, Williams B, Enver T. Increased Complement Receptor-3 levels in monocytes and granulocytes distinguish COVID-19 patients with pneumonia from those with mild symptoms. International Journal of Infectious Diseases. 2020.

10. Herrmann JB, Muenstermann M, Strobel L, Schubert-Unkmeir A, Woodruff TM, Gray-Owen SD, et al. Complement C5a Receptor 1 Exacerbates the Pathophysiology of N. meningitidis Sepsis and Is a Potential Target for Disease Treatment. mBio. 2018;9(1):e01755–17.

11. Mollnes TE, Brekke OL, Fung M, Fure H, Christiansen D, Bergseth G, et al. Essential role of the C5a receptor in E coli-induced oxidative burst and phagocytosis revealed by a novel lepirudin-based human whole blood model of inflammation. Blood. 2002;100(5):1869–77.

12. Pfister F, Vonbrunn E, Ries T, Jäck H-M, Überla K, Lochnit G, et al. Complement Activation in Kidneys of Patients With COVID-19. Frontiers in immunology. 2021;11(3833).

13. Yan B, Freiwald T, Chauss D, Wang L, West E, Mirabelli C, et al. SARS-CoV-2 drives JAK1/2-dependent local complement hyperactivation. Science Immunology. 2021;6(58).

14. Boussier J, Yatim N, Marchal A, Hadjadj J, Charbit B, El Sissy C, et al. Severe COVID-19 is associated with hyperactivation of the alternative complement pathway. The Journal of allergy and clinical immunology. 2021.

15. Valenti L, Griffini S, Lamorte G, Grovetti E, Uceda Renteria SC, Malvestiti F, et al. Chromosome 3 cluster rs11385942 variant links complement activation with severe COVID-19. Journal of Autoimmunity. 2021;117:102595.

16. Eriksson O, Hultström M, Persson B, Lipcsey M, Ekdahl KN, Nilsson B, et al. Mannose-Binding Lectin is Associated with Thrombosis and Coagulopathy in Critically Ill COVID-19 Patients. Thrombosis and haemostasis. 2020.

17. Ramlall V, Thangaraj PM, Meydan C, Foox J, Butler D, Kim J, et al. Immune complement and coagulation dysfunction in adverse outcomes of SARS-CoV-2 infection. Nature medicine. 2020.

18. Jarlhelt I, Nielsen SK, Jahn CXH, Hansen CB, Pérez-Alós L, Rosbjerg A, et al. SARS-CoV-2 Antibodies Mediate Complement and Cellular Driven Inflammation. Frontiers in immunology. 2021;12(4612).

19. Charitos P, Heijnen I, Egli A, Bassetti S, Trendelenburg M, Osthoff M. Functional Activity of the Complement System in Hospitalized COVID-19 Patients: A Prospective Cohort Study. Frontiers in immunology. 2021;12:765330.

20. Defendi F, Leroy C, Epaulard O, Clavarino G, Vilotitch A, Le Marechal M, et al. Complement Alternative and Mannose-Binding Lectin Pathway Activation Is Associated With COVID-19 Mortality. Frontiers in immunology. 2021;12:742446.

21. Yu J, Yuan X, Chen H, Chaturvedi S, Braunstein EM, Brodsky RA. Direct activation of the alternative complement pathway by SARS-CoV-2 spike proteins is blocked by factor D inhibition. Blood. 2020.

22. Ali YM, Ferrari M, Lynch NJ, Yaseen S, Dudler T, Gragerov S, et al. Lectin Pathway Mediates Complement Activation by SARS-CoV-2 Proteins. Frontiers in immunology. 2021;12:714511.

23. Kocsis A, Kekesi KA, Szasz R, Vegh BM, Balczer J, Dobo J, et al. Selective inhibition of the lectin pathway of complement with phage display selected peptides against mannose-binding lectin-associated serine protease (MASP)-1 and -2: significant contribution of MASP-1 to lectin pathway activation. Journal of immunology (Baltimore, Md : 1950). 2010;185(7):4169–78.

24. Chhabra M, Wimmer N, He QQ, Ferro V. Development of Improved Synthetic Routes to Pixatimod (PG545), a Sulfated Oligosaccharide-Steroid Conjugate. Bioconjugate Chemistry. 2021;32(11):2420–31.

25. Li XX, Lee JD, Massey NL, Guan C, Robertson AAB, Clark RJ, et al. Pharmacological characterisation of small molecule C5aR1 inhibitors in human cells reveals biased activities for signalling and function. Biochemical pharmacology. 2020;180:114156.

26. Brekke OL, Christiansen D, Fure H, Fung M, Mollnes TE. The role of complement C3 opsonization, C5a receptor, and CD14 in E. coli-induced up-regulation of granulocyte and monocyte CD11b/CD18 (CR3), phagocytosis, and oxidative burst in human whole blood. Journal of leukocyte biology. 2007;81(6):1404–13.

27. Zheng S, Fan J, Yu F, Feng B, Lou B, Zou Q, et al. Viral load dynamics and disease severity in patients infected with SARS-CoV-2 in Zhejiang province, China, January-March 2020: retrospective cohort study. BMJ. 2020;369:m1443.

28. Tsukagoshi H, Shinoda D, Saito M, Okayama K, Sada M, Kimura H, et al. Relationships between Viral Load and the Clinical Course of COVID-19. Viruses. 2021;13(2).

29. Loeven MA, Rops AL, Berden JH, Daha MR, Rabelink TJ, van der Vlag J. The role of heparan sulfate as determining pathogenic factor in complement factor H-associated diseases. Molecular immunology. 2015;63(2):203–8.

30. Clausen TM, Sandoval DR, Spliid CB, Pihl J, Perrett HR, Painter CD, et al. SARS-CoV-2 Infection Depends on Cellular Heparan Sulfate and ACE2. Cell. 2020;183(4):1043-57.e15.

31. Sarrazin S, Lamanna WC, Esko JD. Heparan sulfate proteoglycans. Cold Spring Harbor Perspectives in Biology. 2011;3(7).

32. Zheng Y, Zhao J, Li J, Guo Z, Sheng J, Ye X, et al. SARS-CoV-2 spike protein causes blood coagulation and thrombosis by competitive binding to heparan sulfate. International Journal of Biological Macromolecules. 2021;193:1124–9.

33. Amarilla AA, Modhiran N, Setoh YX, Peng NYG, Sng JDJ, Liang B, et al. An Optimized High-Throughput Immuno-Plaque Assay for SARS-CoV-2. Frontiers in microbiology. 2021;12(75).

34. Brekke OL, Hellerud BC, Christiansen D, Fure H, Castellheim A, Nielsen EW, et al. Neisseria meningitidis and Escherichia coli are protected from leukocyte phagocytosis by binding to erythrocyte complement receptor 1 in human blood. Molecular immunology. 2011;48(15-16):2159–69.

35. Watterson D, Wijesundara DK, Modhiran N, Mordant FL, Li Z, Avumegah MS, et al. Preclinical development of a molecular clamp-stabilised subunit vaccine for severe acute respiratory syndrome coronavirus 2. Clinical & Translational Immunology. 2021;10(4):e1269.

